# Variable PD-1 glycosylation modulates the activity of immune checkpoint inhibitors

**DOI:** 10.1101/2023.08.25.554811

**Authors:** Chih-Wei Chu, Tomislav Čaval, Frederico Alisson-Silva, Akshaya Tankasala, Christina Guerrier, Gregg Czerwieniec, Heinz Läubli, Flavio Schwarz

## Abstract

Monoclonal antibodies targeting the immune checkpoint PD-1 have provided significant clinical benefit across a number of solid tumors, with differences in efficacy and toxicity profiles possibly related to their intrinsic molecular properties. Here, we report that camrelizumab and cemiplimab engage PD-1 through interactions with its fucosylated glycan. Using a combination of protein and cell glycoengineering, we demonstrate that the two antibodies bind preferentially to PD-1 with a core fucose at the asparagine N58 residue. We then provide evidence that the concentration of fucosylated PD-1 in the blood of non-small cell lung cancer patients varies across different stages of disease. This study illustrates how glycoprofiling of surface receptors and related circulating forms can inform the development of differentiated antibodies that discriminate glycosylation variants and achieve enhanced selectivity and paves the way towards the implementation of personalized therapeutic approaches.

## Introduction

PD-1 is a type-I transmembrane glycoprotein expressed in immune cells, predominantly in T cells, that functions as a checkpoint: engagement of PD-1 by its ligands PD-L1 and PD-L2 suppresses T cell receptor (TCR)-driven cell activation and induces apoptosis^1^. This mechanism can be detrimental in the tumor microenvironment, where a combination of exhausted T cells displaying PD-1 and PD-L1 expression by cancer and myeloid cells cause ineffective antitumor immune responses^2^. Consequently, monoclonal antibodies that target PD-1 or its ligands and prevent their interaction have been developed as anticancer therapeutics^3–6^. Whereas immune checkpoint inhibitors have revolutionized cancer treatment across many tumor indications and demonstrated the usefulness of interjecting the PD-1:PD-L1 axis, only a fraction of patients benefit from these drugs and develop durable clinical responses^7,8^. This suggests that critical information about the PD-1 pathway and the mechanisms of signaling blockade may still be unavailable. Moreover, the intrinsic properties of anti-PD-1 antibodies might be related to differences in efficacy or safety observed in the clinical settings.

PD-1 and PD-L1 are regulated both at the transcriptional and post-translational level. PD-1 is rapidly induced upon TCR-mediated activation, decreases with antigen clearance, but it’s maintained on antigen-specific T cells in chronic settings as cancer^9^. PD-1 stability at the cell surface is controlled by ubiquitination that leads to protein degradation^10^ and by glycosylation that may occur at four asparagine sites (N49, N58, N74 and N119). CRISPR-based screening identified Fut8, which encodes for a fucosyltransferase that adds alpha1,6 core fucose to N-glycans, as a mechanism that regulates cell surface expression of PD-1 by modification of glycans at sites N49 and N74^11^. In line with this observation, T cells exposed to a fucose inhibitor produce stronger antitumor responses^11,12^. Similarly, PD-L1 expression and stability are regulated at the post-translational level by glycosylation and ubiquitination^13^. PD-L1 glycosylation is also critical for interaction with PD-1; in contrast, PD-1 glycosylation appears to be dispensable for binding to PD-L1 as its glycosylation sites are distant from the ligand binding interface^10^. Interestingly, some antibodies, including cemiplimab and camrelizumab, interact with a region surrounding the N58 glycosylation site and require the N58 glycan for efficient binding^14,15^. This property is not shared by other antibodies, such as nivolumab and pembrolizumab, that have partly overlapping binding epitopes interacting with the flexible N- and C’D-loops of the PD-1 molecule, respectively^16–19^. In addition to the transmembrane forms, soluble PD-1 and PD-L1 variants (sPD-1 and sPD-L1) can be generated by protease-based cleavage and accumulate in blood^20,21^. While the function of the released protein variants is not fully established, it has been reported that sPD-L1 retains its inhibitory capacity^22^ and elevated sPD-L1 levels have been associated with advanced disease and worse prognosis^23^. sPD-1 appears to be less informative as a prognostic or predictive biomarker^20^. Notably, the glycosylation status of sPD-1 or sPD-L1 has not been determined.

Here, we observed that camrelizumab and cemiplimab specifically interact with a fucose moiety within the N58 glycan of PD-1. As fucosylation is increased in cancer and fucosylated biomarkers in blood have been associated with a lack of benefit of immune checkpoint inhibitor therapy^24,25^, we investigated the influence of fucosylation on PD-1 in antibody activity by a combination of protein- and cell-based assays. Furthermore, we characterized fucosylation of sPD-1 in serum of individuals with non-small cell lung cancer (NSCLC). These data illustrate a path towards the development of personalized immunotherapeutic approaches for cancer treatment and highlight the potential of glycosylation analysis of targets to guide the development of differentiated therapeutics.

## Materials and methods

### Reagents

2-Fluoro-peracetyl-fucose (2FPF) was purchased from Cayman Chemicals (Ann Arbor, MI) and dissolved in DMSO (Cell Signaling Technologies, Danvers, MA). The anti-human PD-1 antibodies camrelizumab, pembrolizumab, nivolumab and cemiplimab were from Selleck Chemicals (Houston, TX). Human IgG4 isotype was from Biolegend (San Diego, CA).

### Serum samples

Pre-treatment NSCLC serum samples were sourced from iSpecimen (Lexington, MA). Samples were collected in accordance with relevant applicable guidelines and regulations for human subjects’ protection as outlined in the Declaration of Helsinki. All protocols were approved under respective Institutional Review Boards and Ethics Committees (20223899). All subjects utilized in this study provided a written informed consent prior to collection of samples. Collected clinical data included age at blood draw, body mass index, sex, race, and histopathological data and clinical staging, when applicable.

### Cell lines and culture

CHO-K1 and CHO-K1 Fut8 KO cell lines (Creative Biogene, Shirley, NY) were cultured in RPMI-1640 media (Thermo Fisher Scientific, Walham, MA) supplemented with 10% fetal bovine serum (FBS) and 1% penicillin-streptomycin. Jurkat-Lucia TCR-hPD-1 and Raji-APC-hPD-L1 cells were purchased from InvivoGen (San Diego, CA) and cultured in IMDM media (Thermo Fisher Scientific) supplemented with 10% fetal bovine serum (VWR, Radnor, PA), 1% penicillin-streptomycin (Thermo Fisher Scientific) and 100 ug/ml Normocin (Invivogen). Adherent cell lines were maintained in a humidified incubator at 37°C with 5% CO2. Expi293 cells (Thermo Fisher Scientific) were maintained in Expi293 expression media (Thermo Fisher Scientific) in a humidified incubator at 37°C with 8% CO2 at 125 rpm.

### Recombinant protein production

DNA encoding for residues 24-167 of the extracellular portion of human PD-1 (UniProt Q15116) with a C-terminus histidine tag or corresponding PD-1 N58Q mutant were cloned into pcDNA3.4 by GenScript (Piscataway, NJ). Expi293 cells were transiently transfected with plasmid DNA mixed with PEI (Polysciences, Warrington, PA) and incubated for 5 days at 37 degC. For expression of fucosylated PD-1, Expi293 cells were grown in presence of 0.6 mM 2FPF. PD-1 proteins were purified by affinity chromatography using Ni Sepharose Excel resin (Cytiva, Marlborough, MA). PD-10 columns (Cytiva) were used to exchange buffer to phosphate-buffered saline (PBS; Corning, Tweksbury, MA). Protein quality and purity was assessed by SDS-PAGE (BioRad, Hercules, CA) and size-exclusion chromatography using a Superdex 200 5/150 (Cytiva) with Acquity UPLC (Waters, Milford,MA). Protein concentration was determined using a NanoDrop (Wilmington, DE).

### Mass spectrometry analysis of recombinant PD-1 variants

For chymotrypsin digestion, 5 µg of each PD-1 variant were diluted with 50 mM ammonium bicarbonate buffer (Sigma Aldrich) and reduced with the addition of 5 mM dithiothreitol (DTT; Sigma Aldrich). Samples were incubated at 60 ℃ for 20 minutes and then alkylated with 10 mM iodoacetamide (Sigma Aldrich) in the dark at 25℃ for one hour. Sequencing grade chymotrypsin (Promega, Madison, WI) was added at a protein:enzyme ratio of 40:1 and the samples were incubated at 25℃ for 8 hours. For trypsin digestion, 5 µg of each PD-1 variant were diluted in a 50 mM ammonium bicarbonate buffer (Sigma Aldrich) containing 5 mM DTT concentration. Samples were incubated at 60℃ for 45 minutes and then alkylated with 10 mM iodoacetamide for 30 minutes at 25℃. MS-grade trypsin (Pierce) was added to the samples at a protein:enzyme ratio of 40:1. The samples were then incubated at 37℃ for 8 hours before LC-MS grade formic acid (LiChroPur) was added to a final concentration of 1% to quench the reaction.

Digested samples were analyzed using an Orbitrap Exploris 480 Mass Spectrometer (Thermo Scientific) after desalting with an Acclaim Pepmap 100 C18 trap (Thermo Scientific) and liquid chromatography separation with an Ultimate 3000 RSLCnano System (Dionex). Samples eluting from the LC were ionized with a Nanospray Flex™ Ion Source at a spray voltage of 2800V. Each sample was acquired using a Top 20 data dependent acquisition (DDA) method. Raw data were analyzed with Glyco-Decipher software suite^22^ with trypsin or chymotrypsin set as an protease, depending on the experiment, allowing up to 3 missed cleavages. Cysteine carbamidomethylation was set as a fixed modification, and methionine oxidation as variable. Spectrum expansion was enabled, and the other settings were left as default. GlycoTouCan^23^ was used as a glycan database. Identified spectra were manually validated for correct sequence and glycan assignment, quantitation was performed based on summed peak areas of the elution profiles of each glycopeptide at all the detected charge states.

### Antibody binding assays

Binding kinetics of PD-1 antibodies were determined using an Octet Red96 instrument (ForteBio), following previously described procedures^26^. All proteins were diluted in Octet kinetics buffer (Sartorius, Göttingen, Germany). Antibodies were captured on anti-human Fc AHC2 tips (Sartorius). Data capture was performed using Octet Data Acquisition software version 9.0 (ForteBio). Data analysis was performed using Octet Data Analysis software version 9.0 (ForteBio) using a 1:1 binding model.

Antibody binding was also assayed by flow cytometry using CHO-K1 and CHO-K1 *Fut8* KO cells transfected with a human PD-1 plasmid (GenScript) by lipofectamine (Thermo Fisher Scientific). Anti-PD-1 antibodies or human IgG4 isotype were added to the cells and incubated for 20 minutes on ice. Cells were washed and stained with an 0.1 mg/l anti-human IgG Fc-PE antibody for 20 minutes. Cells were analyzed using a CytoFlex instrument (Beckman Coulter, Brea, CA) and data were analyzed using FlowJo software (FlowJo, Ashland, OR).

Additionally, antibody binding was assessed by flow cytometry using human naive CD8 T cells. Briefly, 2 million CD8 T cells (Stemcell Technologies, Cambridge, MA) from three donors were cultured for 48 hours in AIM V Medium (ThermoFisher Scientific) supplemented with 5% heat-inactivated fetal bovine serum and activated with Dynabeads Human T-Activator CD3/CD28 (ThermoFisher Scientific) at a 1:1 bead:cell ratio according to the manufacturer’s instructions, in presence or absence of 0.1 mM of 2FPF. At days 1 and 2 after stimulation, cells were stained with a Zombie Red viability dye kit (Biolegend), followed by fixation with fixation solution (eBioscience-Thermo). Core fucosylation was assessed by incubating cells with FITC-conjugated LCA lectin (Vector Labs) at a 1:500 dilution. To confirm activation followed by bead stimulation, cells were stained with BV711-conjugated anti-CD25 antibody (Clone M-A251, Biolegend) at a 1:50 dilution in PBS. Cells were incubated with pembrolizumab, camrelizumab or human IgG4 isotype control (at 1 nM concentration), followed by incubation with PE-conjugated anti-human IgG4 (Southern Biotechnologies) at a 1:300 dilution in PBS. Cells were analyzed by a CytoFlex flow cytometer and data analysis was performed using FlowJo 10.9.0 software.

### Ligand blocking assays

To assess the binding of PD-L1 to PD-1, biotinylated human PD-L1 (ACROBioSystems, Newark, DE) and PE-conjugated streptavidin (Biolegend, San Diego, CA) were mixed at a 1:2 molar ratio, added to PD-1 transfected CHO-K1 or CHO-K1 *Fut8* KO cells and incubated for 20 minutes on ice. Cells were washed and analyzed by a CytoFlex flow cytometer. To investigate blocking of PD-L1 binding, PD-1 transfected CHO-K1 and CHO-K1 *Fut8* KO cells were incubated with antibodies for 15 minutes on ice before addition of PD-L1:SA-PE complexes. Cells were then washed and analyzed by a CytoFlex flow cytometer.

For plate-based ligand blocking assay, 96-well flat-bottom plates (Thermo Fisher Scientific) were coated with 5 ug/ml of purified recombinant PD-1, PD-1 NF or PD-1 N58Q in PBS overnight at room temperature. Plates were then washed with PBS containing 0.05% Tween 20 (PBST). After blocking with 5% BSA in PBS for one hour at room temperature, wells were washed with PBST, followed by incubation with anti-PD-1 antibodies diluted in PBS containing 0.5% BSA for one hour incubation at room temperature. After washing with PBST, 1ug/ml PD-L1-Fc-biotin in PBS containing 0.5% BSA were added to each well and incubated for two hours. After washing with PBST, the wells were incubated with 0.1 ug/ml horseradish peroxidase-conjugated streptavidin (ACROBiosystems) for 30 minutes. Wells were washed and incubated with 3,3′,5,5′-Tetra-methyl benzidine substrate (Biolegend, San Diego, CA), followed by a stop solution (Biolegend). Absorbance at 450 nm was read with the Envision microplate reader (Perkin Elmer).

### PD-1:PD-L1 functional assay

Jurkat TCR-hPD-1 cells were pre-cultured with or without 300 uM 2F-Peracetyl-Fucose for 3 days in IMDM medium supplemented with 10% FBS, 1% penicillin-streptomycin. Antibodies were diluted in PBS and added into 96 well flat-bottom plates. Jurkat-Lucia TCR-hPD-1 cells and Raji-APC-hPD-L1 were added to the wells and plates were incubated at 37°C in a CO2 incubator for 6 hours. Supernatants were transferred into a 96-well white plate (VWR) and assayed with QUANTI-Luc 4 reagent (Invivogen). Luminescence was read immediately with an Envision system.

### Serum PD-1 quantification

Serum and recombinant PD-1 was quantified by human PD-1 Quantikine ELISA and human PD-1 DuoSet ELISA (R&D Systems, Minneapolis, MN), following manufacturer’s protocol. In addition, serum and recombinant PD-1 were quantified by ELISA assay using wells coated overnight with 5 ug/ml camrelizumab or pembrolizumab in PBS, followed by incubation with 1% BSA in PBS and detection with human PD-1 DuoSet ELISA detection antibody. Wells were washed and incubated with 3,3′,5,5′-Tetra-methyl benzidine substrate. Reaction was stopped with acid solution, and absorbance was read with the Envision microplate reader.

### Data analysis and visualization

GraphPad Prism (GraphPad Software, Boston, MA) was used to represent data and for statistical analysis.

### Protein structure analysis

Interactions and interfaces between antibodies or ligands and PD-1 were calculated with PDBePISA (https://www.ebi.ac.uk/pdbe/pisa/papers/pisa-web.pdf) and CoCoMaps https://www.molnac.unisa.it/BioTools/cocomaps/ using PDB structures 7CU5, 7WVM, 7ZQK, 5WT9 and 5DK3). Structures were visualized by PyMol (Schroedinger, New York, NY).

## Results

### Glycosylation-dependent anti-PD-1 antibodies interact with the core fucose

A number of monoclonal antibodies have been approved as anti-cancer treatment in a broad range of indications (**Suppl Table 1**) with clinical efficacy and toxicity profiles that may be related to the intrinsic properties of the molecule. A subgroup of these antibodies, including camrelizumab and cemiplimab, have been recently identified for their dependency on the N58 glycosylation for binding^14,15^. To elucidate the detailed binding characteristics of camrelizumab and cemiplimab to PD-1 and the PD-L1 mechanism, we inspected the available structures of antibody:PD-1 complexes. In line with what was previously described, we observed that camrelizumab and cemiplimab bound to a similar region of PD-1 in proximity of the N58 glycosylation site, whereas pembrolizumab and nivolumab recognized different epitopes (**Suppl Fig 1**). In particular, the presence of the N58 glycan of PD-1 in the camrelizumab:PD-1 complex allowed us to visualize the interactions between the VH and the glycan. In addition to the previously described interactions between the HCDR1 and HCDR2 and the core glycan, we observed that residues S30 and S31 of the CRD1 and S52, G53 and G54 of the CRD2 engaged with the core alpha1,6 fucose through hydrogen bonds (**Figure 1**). Although the PD-1 N58 glycan was not visible in the complex with cemiplimab, the antibody appeared to bind to PD-1 in a similar orientation as camrelizumab, suggesting the N58 glycan composition may influence binding and activity of both antibodies.

**Figure 1.**
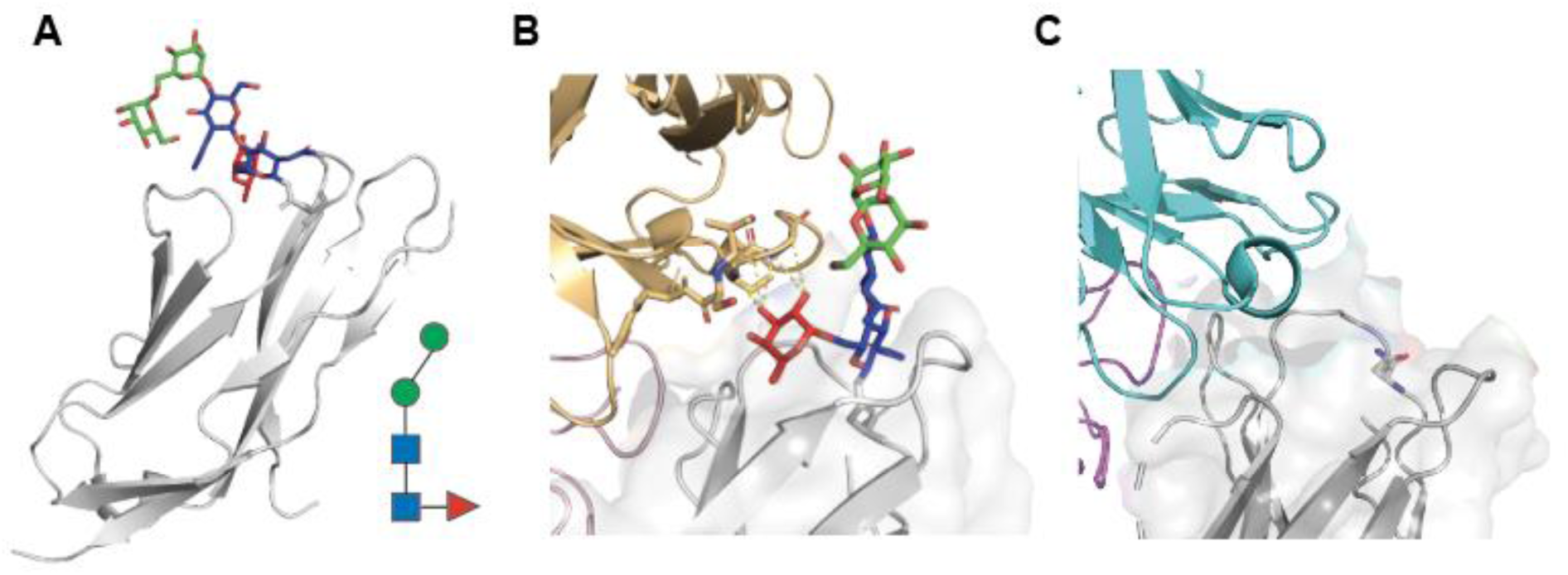
The binding interface between PD-1 and the anti-PD-1 antibodies camrelizumab and cemiplimab includes the core fucose of the PD-1 N58 glycan. **A.** Structure of PD-1 (grey) with the glycan linked to the asparagine N58 of the BC loop of PD-1. The glycan visible in the crystal structure (PDB 7CU5) includes a fucose, two *N-*acetyl-glucosamine units and two mannose units. The glycan structure is also represented with geometric symbols as convention (red triangle: fucose; blue square: *N-*acetyl-glucosamine; green circle: mannose). **B**. Structure of PD-1 in complex with camrelizumab (PDB 7CU5) indicating potential interactions between the core fucose of PD-1 N58 glycan and the heavy chain of the antibody (residues S30 and S31 of CRD1 and S52, G53 and G54 of CRD2) The VH chain is indicated in gold and the VL is in pink. **C**. Structure of cemiplimab in complex with PD-1 (PDB 7WVM). The N58 residue of PD-1 is represented as sticks. Cemiplimab binds to PD-1 in a similar orientation as camrelizumab, suggesting the N58 glycan structure may influence affinity. VH is in cyan and VL is in purple.

### Core fucose of the PD-1 N58 glycan is a key determinant for camrelizumab and cemiplimab binding

To directly test if the core fucose of the PD-1 N58 glycan was involved in antibody binding, we used both protein- and cell-based assays. First, we generated three recombinant PD-1 variants by expressing the extracellular region of PD-1 in Expi293 cells in presence or absence of the fucose inhibitor 2-fluoro-peracetyl-fucose, and by producing a N58Q mutant protein (**Fig. 2A**). Mass spectrometry analysis confirmed that the glycoforms at the N58 site of the PD-1 and PD-1 NF (expressed in presence of fucose inhibitor) variants were comparable and differed only for the presence or absence of fucose (**Fig. 2B** and **Supplementary Fig. 2-4**). Binding of camrelizumab and cemiplimab to the three PD-1 variants was examined by a biolayer interferometry assay, along with nivolumab and pembrolizumab as controls. We observed a 100-fold difference in affinity of camrelizumab to PD-1 and PD-1 N58Q, in line with previous reports (**Table 1**). Strikingly, the difference in KD between PD-1 and the N58Q mutant was completely recapitulated by PD-1 NF, indicating that the contribution of the N58 glycan to the antibody binding is largely provided by the core fucose. Similarly, binding of cemiplimab to PD-1 was compromised by the absence of fucose, although not to the same extent as camrelizumab. As expected, binding of pembrolizumab or nivolumab to the PD-1 variants was not affected by changes in glycosylation.

**Figure 2.**
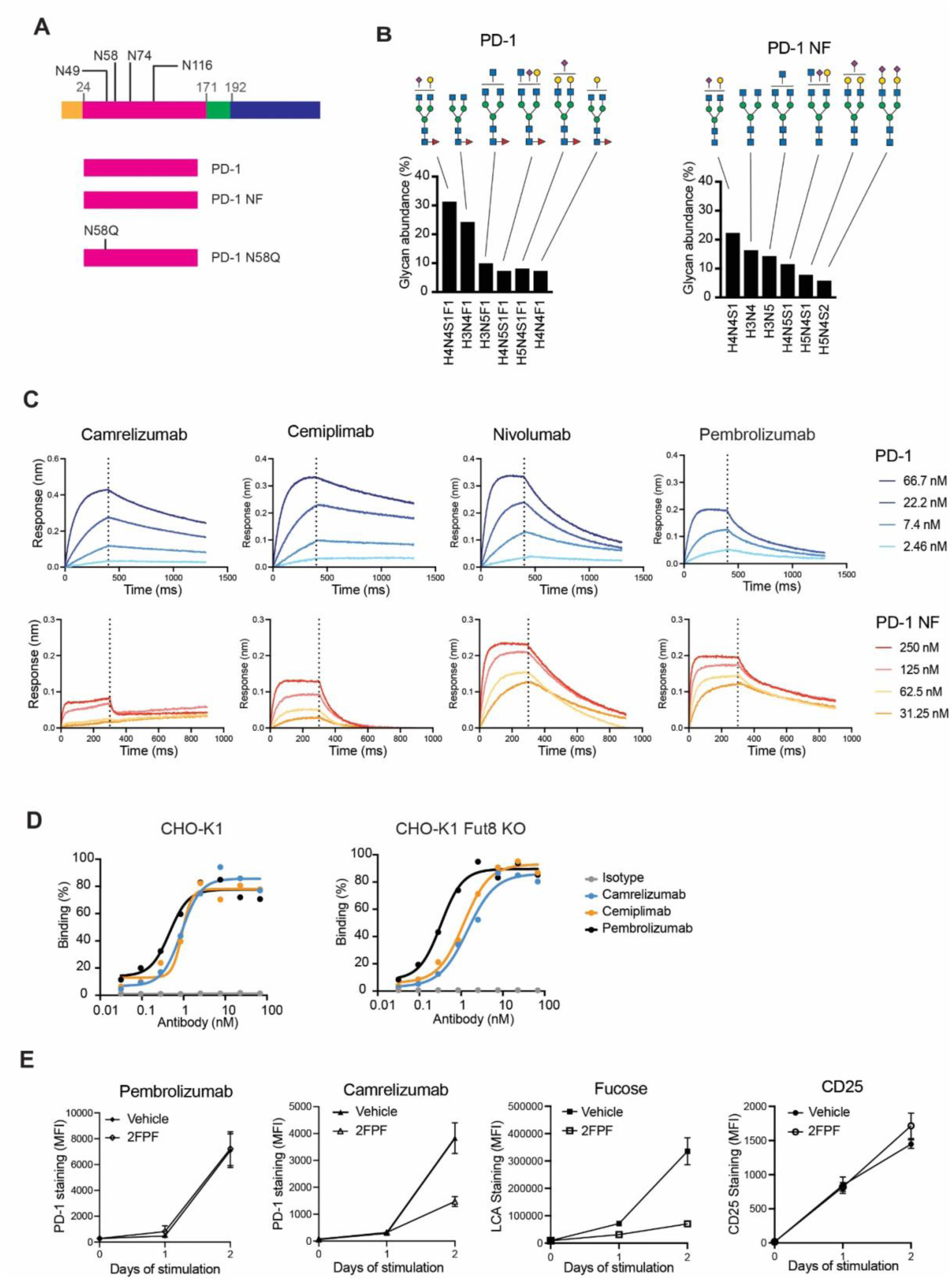
Binding of camrelizumab and cemiplimab to PD-1 depends on the core fucose of the N58 glycan of PD-1. **A.** Schematic diagram of the PD-1 proteins indicating signal peptide (in orange), extracellular domain (in pink), transmembrane domain (in green) and intracellular region (in blue), as well as the four N-linked glycosylation sites. **B.** Relative abundance of the glycoforms found at the N58 site of recombinant PD-1 proteins. Glycan structures are comparable and differ only by the fucose moiety (red triangle). Glycan composition is indicated at the bottom of the graph (H: hexose; N: N-acetyl-hexosamine; S:sialic acid; F: fucose) and represented with symbols at the top. **C.** Representative binding profiles of anti-PD-1 antibodies to recombinant PD-1 proteins. The dotted line marks the separation between the association and dissociation cycle. **D**. Binding of anti-PD-1 antibodies to CHO-K1 cells transfected with full length PD-1 constructs. Lack of fucosyltransferase FUT8 expression that adds core fucose does not alter the binding profile of pembrolizumab but results in reduced binding by cemiplimab and camrelizumab. **E**. Binding of anti-PD-1 antibodies to human CD8 T cells. Cells were activated with CD3/CD28 beads in presence or absence of fucose inhibitor and tested for expression of PD-1, fucose or CD25. Detection of PD-1 by camrelizumab depends on fucose content, whereas neither pembrolizumab nor the anti-CD25 antibodies are affected by changes in fucosylation. Vehicle refers to the buffer used for 2FPF solubilization.

**Table 1.** Affinity of anti-PD-1 antibodies for PD-1 variants.

|  | KD (M) |  |  |  |
| --- | --- | --- | --- | --- |
|  | Camrelizumab | Cemiplimab | Pembrolizumab | Nivolumab |
| PD-1 | 3.40E-09±5.12E-11 | 2.95E-09±1.64E-10 | 2.25E-09±5.99E-11 | 4.96E-09±1.87E-10 |
| PD-1 NF | 1.90E-07±1.05E-08 | 8.71E-08±1.33E-09 | 3.67E-09±7.35E-11 | 2.08E-08±2.71E-10 |
| PD-1 N58Q | 4.49E-07±4.53E-08 | 2.61E-07±2.45E-08 | 2.96E-09±4.97E-11 | 1.07E-08±1.19E-10 |

Next, to further test the hypothesis that core fucose was a key determinant for binding, we analyzed antibody binding to full length PD-1 expressed in wild type or Fut8-knockout CHO-K1 cells (**Fig 2C**). Absence of core fucose reduced binding of camrelizumab and cemiplimab, but did not alter binding of pembrolizumab, resulting in relative EC50 values of 0.9, 0.9 and 0.5 nM in CHO-K1 cells and 1.4, 1.15 and 0.3 nM in CHO-K1 Fut8-KO cells for camrelizumab, cemiplimab and pembrolizumab, respectively.

Lastly, we assessed PD-1 detection by camrelizumab in primary human CD8 T cells activated in presence or absence of fucose inhibitor. As expected, naive CD8 T cells expressed zero or relatively low levels of PD-1 and CD25 (**Figure 2E**). CD3/CD28 stimulation induced CD25 expression within 24 hours and PD-1 expression within 48 hours. Remarkably, activation in presence of the fucose inhibitor resulted in a substantial decrease of binding of the lectin LCA and PD-1 detection by camrelizumab, but did not affect recognition by pembrolizumab or the CD25 antibody (**Figure 2E**).

Altogether, these data indicate that both fucosylated and non-fucosylated PD-1 can be expressed on the surface of T cells and that binding of camrelizumab and, to a lesser extent, cemiplimab depends on core fucosylation of the N58 glycan of PD-1.

### PD-1 fucosylation affects PD-L1 blocking efficacy of camrelizumab and cemiplimab

Next, we sought to understand if the glycosylation structure of PD-1 had an impact on the ability of anti-PD-1 antibodies to block interactions with PD-L1 (**Fig 3A**). First, we assessed if the fucosylation status of PD-1 modulated interaction with its ligand, and observed no differences in PD-L1 binding to PD-1 in CHO-K1 cells regardless of *Fut8* expression (EC_50_ values were 1.0 nM for CHO-K1 cells and and 1.7 nM CHO-K1 *Fut8* KO cells) (**Supplementary Fig. 6**). However, whereas ligand blocking profiles were overlapping in CHO-K1 cells, camrelizumab and cemiplimab were less effective in preventing PD-L1 interactions in *Fut8*-deficient cells (calculated IC50 values were 5.6 nM camrelizumab, 2.6 nM for cemiplimab and 0.9 nM for pembrolizumab). To further investigate the impact of fucosylation on the blocking efficacy of anti-PD-1 antibodies, we monitored their capacity to impair binding of PD-L1 to plate-coated PD-1 variants (**Fig 3B**). Inhibition of PD-L1 binding to PD-1 was similar for cemiplimab, camrelizumab and pembrolizumab. However, blockade of PD-L1 binding was sensitive to both fucosylation and glycosylation at the N58 site, with calculated IC50 values 1.6 nM for camrelizumab, 0.6 nM for cemiplimab, and 0.5 nM for pembrolizumab for PD-1 NF and 22.3 nM for camrelizumab, 1.6 nM for cemiplimab, and 0.8 nM for pembrolizumab for PD-1 N58Q.

**Figure 3.**
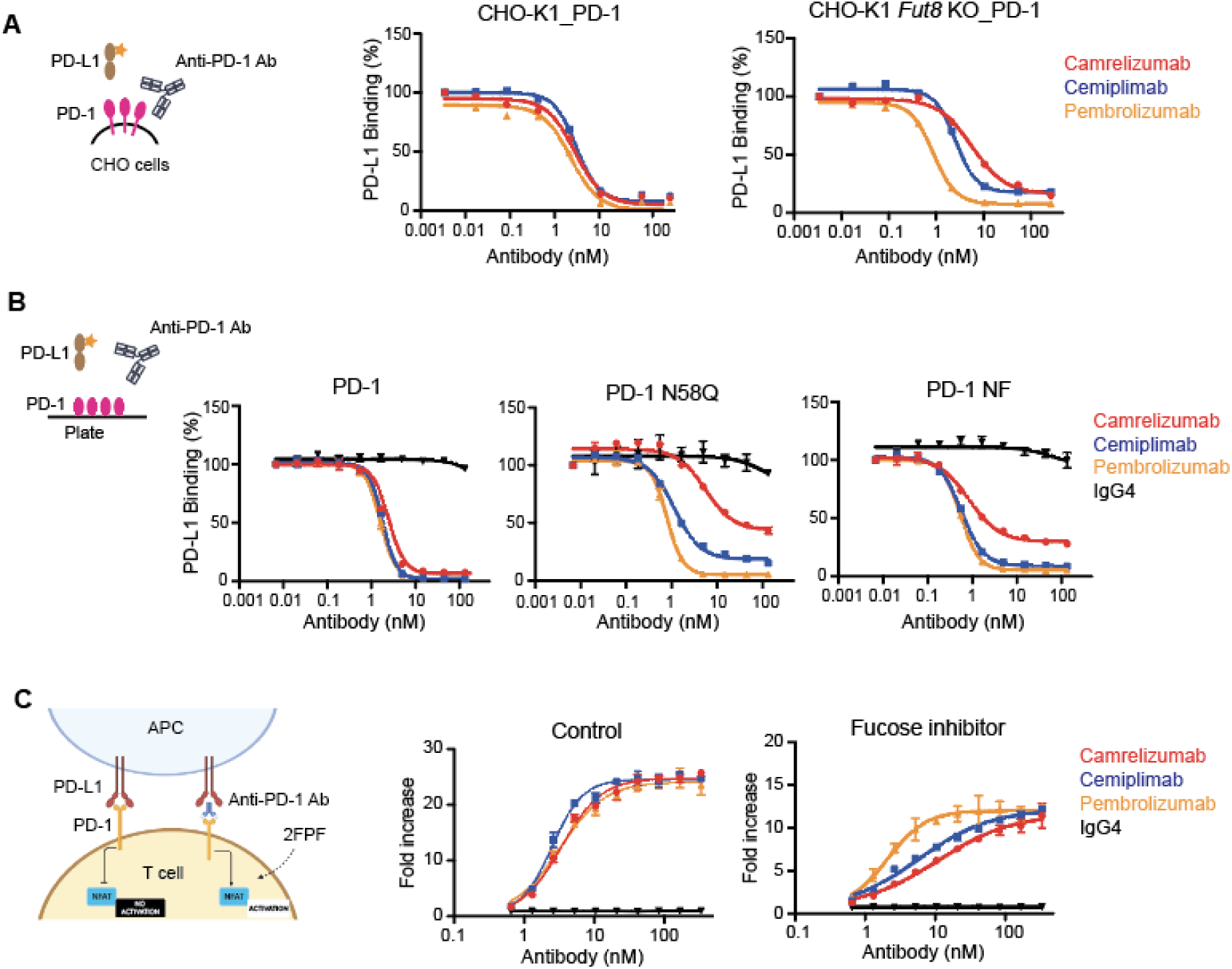
PD-1 Fucosylation impacts the ability of camrelizumab and cemiplimab to block PD-1:PD-L1 interactions. **A.** Blockade of PD-L1 binding to PD-1 expressed in CHO-K1 or fucosyltransferase 8-deficient cells by camrelizumab, cemiplimab or pembrolizumab. PD-L1 Binding was assessed by flow cytometry. Three independent assays were conducted, with each assay performed in duplicate as technical replicates. A representative figure depicting the results is provided.Interpolation was calculated using a 4-parameter logistic function. **B.** Blockade of PD-L1 binding to recombinant by anti-PD-1 antibodies was assessed by a plate-based assay. Three independent assays were conducted, with each assay performed in duplicate as technical replicates. A representative figure depicting the results is provided. **C.** Reduction in fucosylation alters the ability of camrelizumab and cemiplimab to modulate PD-1 signaling. A schematic diagram of the PD-1/PD-L1 cell-based assay, which relies on a reporter T cell line engineered to express PD-1, TCR and CD28, and an antigen-presenting cell line (APC) expressing PD-L1, antigen-MHC complex and CD80. Luciferase activity is used as an indicator of PD-1 signaling activity, which is blocked by PD-L1 on the APC. Luminescence was measured and represented as fold increased over non-induced cells. Three independent assays were conducted, with each assay performed in duplicate as technical replicates. A representative figure depicting the results is provided.

Lastly, to determine if the observed differences in ligand blockade had functional consequences in modulating PD-1 signaling, we employed a reporter assay that uses suppression of luciferase expression as a readout of PD-1:PD-L1 interaction (**Fig 3C**). Consistently with our previous observations, we didn’t detect differences in blocking ability of the anti-PD-1 antibodies in untreated cells. However, treatment of Jurkat hPD-1 cells with a fucose inhibitor (**Suppl. Figure 7**) revealed differences in the potency of the antibodies, with calculated IC50 values of 10.7 nM for camrelizumab, 5.7 nM for cemiplimab, and 2.0 nM for pembrolizumab. The lower overall signals detected in Jurkat cells exposed to the fucose inhibitors might be related to the impact of fucose on other components of the immune synapse. Taken together, these data suggest that the blocking efficacy of the anti-PD-1 antibodies camrelizumab and cemiplimab is influenced by PD-1 fucosylation at the N58 site.

### Fucosylated PD-1 increases in serum of late-stage lung cancer patients

As camrelizumab and cemiplimab are selective for fucosylated PD-1, we sought to understand if there was variability in fucose content of soluble PD-1 in blood of cancer patients. In order to detect fucosylated PD-1, we developed a plate-based assay that employs plate-bound camrelizumab or pembrolizumab for capturing PD-1, followed by detection with polyclonal anti-PD-1 antibodies. The two assays were able to discriminate between fucosylated and non-fucosylated PD-1 (**Suppl Fig 8**). Surprisingly, similar binding patterns were observed with commercial kits (**Suppl Fig 8**). Using combinations of these assays, we were able to measure fucosylated PD-1 in the serum of a cohort of NSCLC patients (**Suppl Table 2**). Whereas there were no differences in the concentration of total serum PD-1 in different disease stages (**Figure 4A**), the fraction of fucosylated PD-1 appeared to increase in late-stage lung cancer (**Figure 4B**). Altogether, these data indicate that there is variability in serum PD-1 fucosylation of cancer patients.

**Figure 4.**
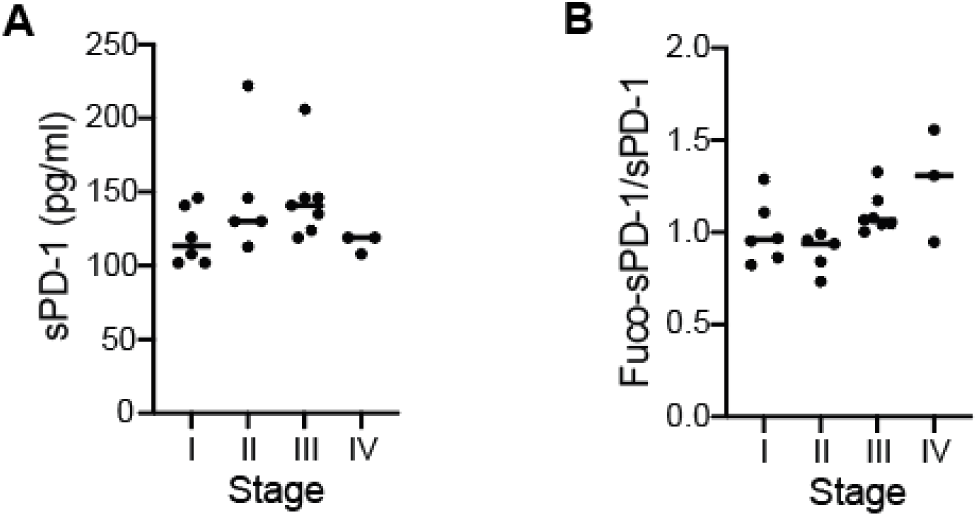
Fucosylation of serum PD-1 of NSCLC patients varies in different disease stages. **A.** Concentration of sPD-1 does not significantly change during progression of disease in this cohort of NSCLC patients. **B.** Fucosylated sPD-1 is higher in serum for late-stage lung cancer patients. The fucosylated sPD-1 fraction was calculated as the ratio of fucosylated sPD-1 divided by total sPD-1 concentration.

## Discussion

Glycosylation impacts the structure of proteins, their activity and interactions with other molecules^27,28^. The availability of increasingly sophisticated tools to characterize this protein modification has allowed the definition of structure-function relationships, bringing light to the complexity of the glycan structures, the fine regulation of the underlying biosynthetic mechanisms, and their detailed roles^29–33^. For example, modulation of IgG1 binding to FcgRIIIA by Fc fucosylation is a well known feature that has been exploited in therapeutics design to enhance antibody-dependent cytotoxicity and antibody-dependent phagocytosis^34^. While currently underexplored, glycoprofiling of new or validated drug targets represents a valuable opportunity for development of next generation therapeutics that may achieve exquisite selectivity by binding to distinct glycosylation variants. Additionally, the recent discovery of fucosylated biomarkers in blood of melanoma patients that failed to achieve extended survival following treatment with immune checkpoint inhibitors point to a role of glycosylation not only in the tumor but also in the periphery^24,25^. Therefore, glycosylation analysis of circulating glycoproteins may offer crucial information for both drug design and biomarker discovery.

In this work, we observed that camrelizumab interacted with the core fucose of the N58 N-glycan of PD-1. We then confirmed that camrelizumab and cemiplimab bound preferentially to PD-1 carrying glycans with core fucose. Camrelizumab exhibited a stronger dependance on this modification compared to cemiplimab, probably due to the contribution of serine S31 in the HCDR1 in binding to the fucose moiety. Using a combination of protein- and cell-based assays, we showed that the ligand blocking activity of these two antibodies is sensitive to variations in fucose. Lastly, we provided preliminary data indicating that fucosylation of soluble PD-1 in blood varies in different stages of lung cancer. Other antibodies that depend on the N58 glycan for binding, including MW11-h317, mAB059c, and STM418, might be regulated in a similar fashion by the glycan structure^35–37^.

Fucosylation can regulate PD-1 function in T cells. Whereas both fucosylated and non fucosylated forms of PD-1 can be found on T cells, the stability of PD-1 may be enhanced by fucosylation at sites N49 and N74^11^. It is interesting that core fucosylation promotes the stability of the cell surface adhesin LCAM1, likely by blocking shedding caused by proteases^38^. Similarly, MHC-II fucosylation accumulates this protein at the surface of tumor cells^39^. Therefore, core fucosylation might exhibit a general role in supporting receptor stability at the cell surface. Interestingly, as fucosylation leads to PD-1 accumulation on the T cell surface and limits activation of murine T cells, it has been suggested that terminally exhausted cells display high levels of fucosylated PD-1. As there is debate around which subsets of exhausted T cells (precursor, or terminally exhausted) can be reactivated by anti-PD-1 antibodies, glycoform-specific antibodies may help provide insights on this matter.

This analysis shows that fucosylation of PD-1 has an impact on binding of clinically used blocking antibodies. In addition to the fucosylation at site N58, it is possible that additional glycans may have an impact on antibody binding. Moreover, whereas we observed a 100-fold difference in KD values between fucosylated and non-fucosylated PD-1 for camrelizumab, it resulted in a 5 to 10-fold difference in PD-1:PD-L1 blocking activity. Additional studies in primary cells with endogenous PD-1 expression are needed to provide a better understanding of the functional relevance of fucosylation. Thirdly, lack of data from clinical trials comparing camrelizumab or cemiplimab to glycosylation-independent anti-PD-1 antibodies does not allow a clear assessment of the additional benefit of targeting fucosylated PD-1. Preclinical and clinical evidence suggests that a significant overall survival across multiple tumor indications is obtained with camrelizumab^40,41^, even if pharmacodynamics data from human phase 1 trials of camrelizumab and nivolumab revealed comparable PD-1 binding affinity (KD equal to 3.31 nM and 3.65 nM, respectively) and receptor occupancy on circulating T lymphocytes (85% and 70%, respectively)^42,43^. Of note, whereas camrelizumab displayed similar rates of grade 3–4 treatment-related adverse events compared to nivolumab and pembrolizumab, ranging 22–25% of the cohorts studied, reactive cutaneous capillary endothelial proliferation was observed much more frequently with camrelizumab compared to nivolumab and pembrolizumab (67.0% vs. 2.4%)^44^. This has been attributed to the unexpected binding to vascular endothelial growth factor receptor 2 (VEGFR2), frizzled class receptor 5 and UL16 binding protein 2 (ULBP2) that may correlate with the side effects of capillary hemangiomas observed in clinical studies with camrelizumab^45^. In summary, we demonstrate that two anti-PD-1 antibodies currently used in clinical routine have a preference for glycosylated PD-1 modified with a fucose, a frequent but variable element in glycoproteins. Future studies should determine PD-1 glycosylation status both in T cells in different activation states and in the blood of patients scheduled to receive immune checkpoint inhibitor therapy. Methods with increased sensitivity, such as glycoproteomics, may prove particularly useful for such analyses. This research supports a framework for the development of antibody therapeutics that target distinct subsets of target proteins. If the glycosylation status of membrane-bound and soluble target is comparable, measuring glycosylation of circulating form provides a convenient strategy to obtain information about the tumor microenvironment. Conversely, if peripheral and tissue glycosylation differ, this feature can be engineered in the antibody specificity to enhance tumor targeting or avoid peripheral sink effects.

## Ethics statement

The studies involving human participants were reviewed and all protocols were approved under respective Institutional Review Boards and Ethics Committees (20223899) The patients/participants provided their written informed consent to participate in this study.

## Author contribution

Conceptualization: HL, FS. Sample acquisition: CG. Data generation and analysis: CC, TC, FAS, AT, GC, FS. Manuscript writing: CC, HL, FS. All authors reviewed and approved the manuscript.

## Acknowledgments

We thank all current and past colleagues at InterVenn for input on the project, and Timothy Ohara for manuscript review. We are grateful to Carlito Lebrilla and Carolyn Bertozzi for scientific discussions and feedback.

