## Supplementary Material for "Variable PD-1 glycosylation modulates the activity of immune checkpoint inhibitors"

### Supplementary Materials

**Supplementary Table 1.** Therapeutic anti-PD-1 antibodies approved or in review in the EU or US. Data adapted from *Antibody therapeutics approved or in regulatory review in the EU or US*. The Antibody Society (2023, May,19)

| International non- proprietary name, US product identifier | Brand name | Target; Format | Indication first approved or reviewed | First EU approval year | First US approval year | Estimated PDUFA date or FDA action |
|---|---|---|---|---|---|---|
| Nivolumab | Opdivo | PD1; Human IgG4 | Melanoma, non-small cell lung cancer | 2015 | 2014 | |
| Pembrolizumab | Keytruda | PD1; Humanized IgG4 | Melanoma | 2015 | 2014 | |
| Cemiplimab, cemiplimab-rwlc | Libtayo | PD-1; Human mAb IgG4 | Cutaneous squamous cell carcinoma | 2019 | 2018 | |
| Dostarlimab, dostarlimab-gxly | Jemperli | PD-1; Humanized IgG4 | Endometrial cancer | 2021 | 2021 | |
| Retifanlimab, retifanlimab-Dlwr | Zynyz™ | PD-1; Humanized IgG4 | Merkel cell carcinoma | In review | 2023 | |
| Serplulimab | HANSIZHUANG | PD-1; Humanized IgG4 | Small cell lung cancer | In review | NA | |
| Camrelizumab | AiRuiKa | PD-1; Humanized IgG4 | Hepatocellular carcinoma | NA | In review | |
| Toripalimab | Tuoyi | PD-1; Humanized IgG4 | Nasopharyngeal carcinoma | In review | In review | December 23, 2022 PDUFA date, but decision was delayed due to travel restrictions. |
| Penpulimab | (Pending) | PD-1; Humanized IgG1 | Metastatic nasopharyngeal carcinoma | NA | In review | Status unknown as of Jan 2023; Real-Time Oncology Review |
| Sintilimab | (Pending) | PD-1; Human IgG4 | Non-small cell lung cancer | NA | In review | CR letter issued Mar 2022 |

| | | | | | | |
|---|---|---|---|---|---|---|
| Tislelizumab | (Pending) | PD-1; Humanized IgG4 | Esophageal squamous cell carcinoma | In review | In review | CR letter issued in Mar 2022 |

**Supplementary Table 2.** Study population characteristics. Counts are followed by the appropriate column-wise percentage, while continuous variables are summarized by medians and standard deviation

| **Characteristics** | **iSpecimen** (N=56) |
|---|---|
| Age | 62.2 (8.7) |
| BMI | 26.6 (3.8) |
| Male | 36 (64.3%) |
| Race | |
| Caucasian | 49 (87.5%) |
| Missing | 7 (12.5%) |
| Disease Category | |
| NSCLC Stage I | 14 (25.0%) |
| NSCLC Stage II | 9 (16.1%) |
| NSCLC Stage III | 18 (32.1%) |
| NSCLC Stage IV | 6 (10.7%) |
| Recurrent NSCLC | 2 (3.6%) |
| Healthy | 7 (12.5%) |
| Histology | |
| Adenocarcinoma | 22 (39.3%) |
| Bronchioloalveolar carcinoma | 1 (1.8%) |
| Mucinous adenocarcinoma | 2 (3.6%) |
| Squamous cell carcinoma | 24 (42.9%) |
| Not applicable | 7 (12.5%) |
| Smoking History | |
| Non-smokers | 16 (28.6%) |
| Former smokers | 2 (3.6%) |
| Smokers | 31 (55.4%) |
| Missing | 7 (12.5%) |

**Supplementary Figure 1. Relevance of PD-1 N58 in the PD-L1 binding mechanism and the epitopes of the PD-1 antibodies. A**. The binding surface of PD-L1 on PD-1 (PDB 4ZQK) is highlighted in green. **B**. The epitopes of PD-1 antibodies (Camrelizumab: PDB 7CU5; Cemiplimab: 7WVM; Nivolumab: 5WT9; Pembrolizumab: 5B8C) are highlighted in color. The N58 residue is represented as sticks and included in a circle.

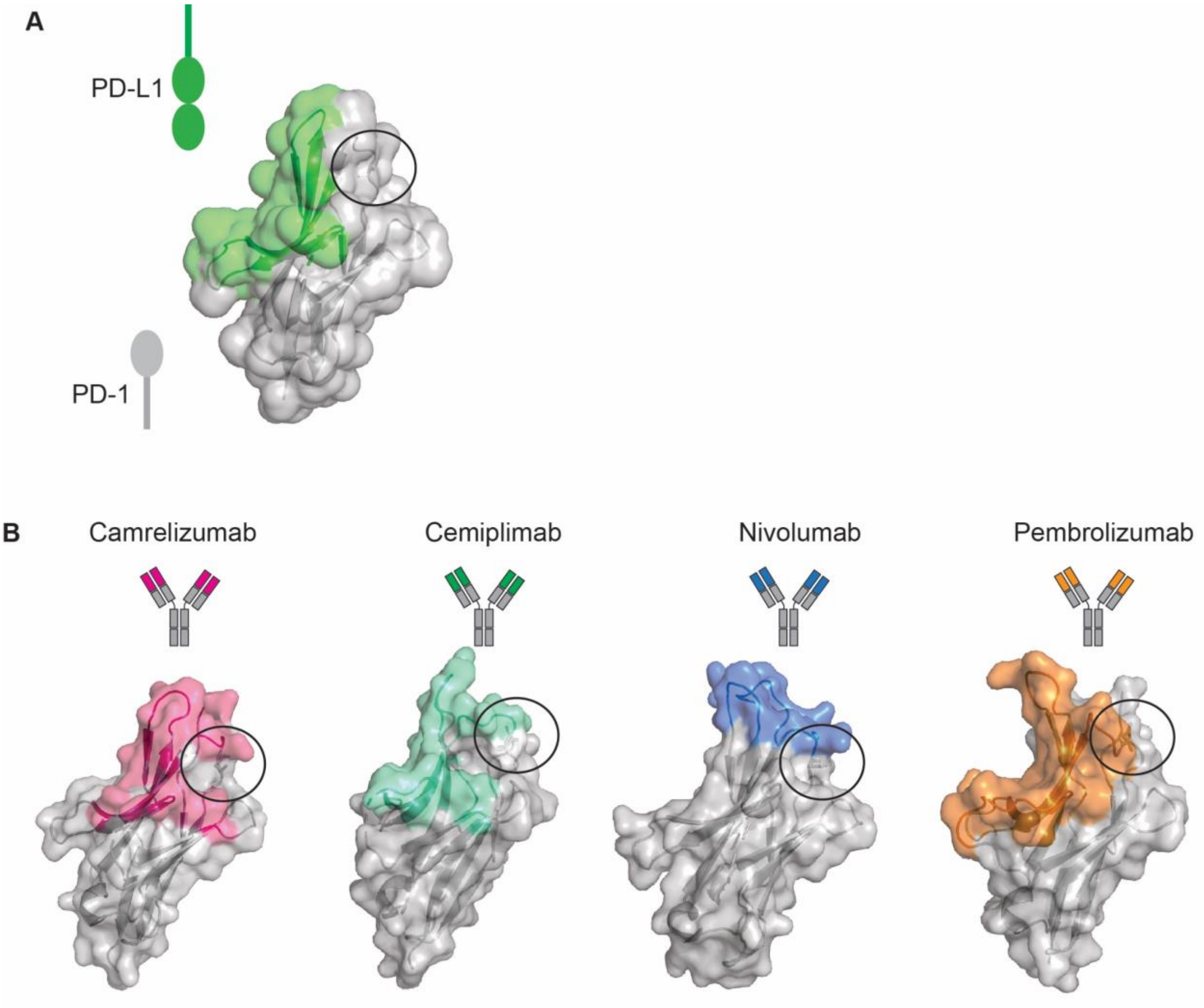

**Supplementary Figure 2. Recombinant expression of PD-1 in presence of fucose inhibitors does not alter overall glycan composition of recombinant PD-1 variant at the three sites N49, N58 and N74.** Glycans are grouped as truncated, oligo-mannose, complex-hybrid and those not found in GlycoTouCan database (Modified).

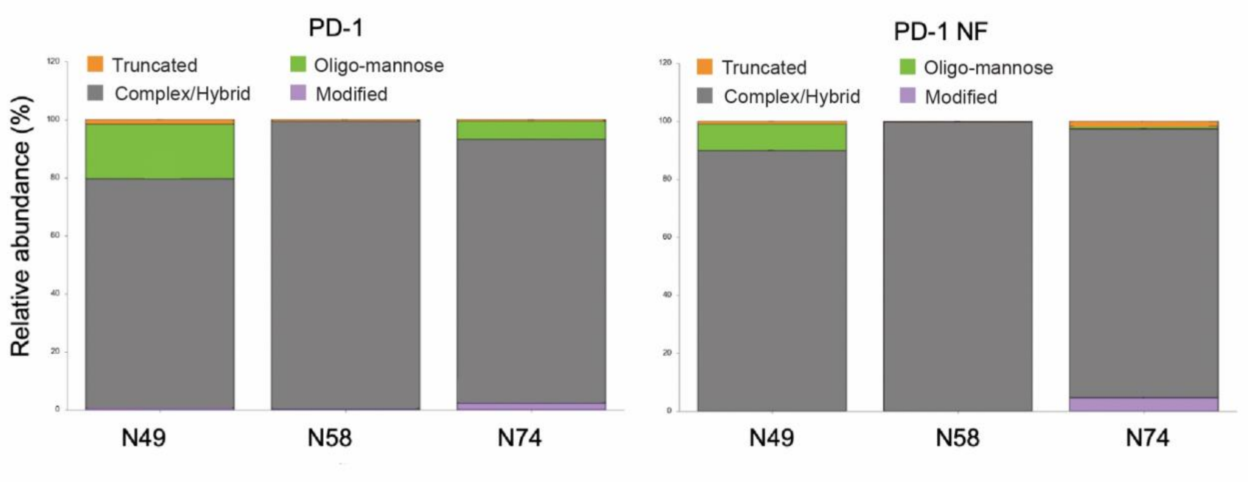

**Supplementary Figure 3. Recombinant expression of PD-1 in presence of fucose inhibitors results in a drastic reduction of fucose content.**

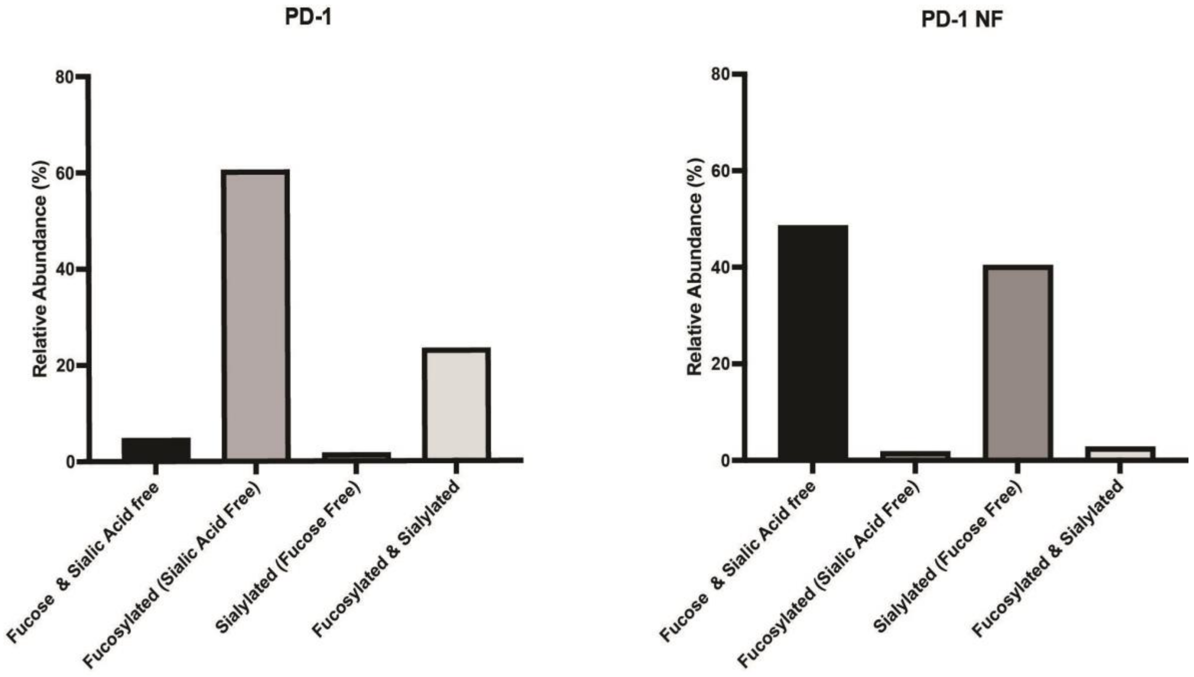

**Supplementary Figure 4. Mass spectrometry analysis of the glycan structures of recombinant PD-1 variants.** The 10 most abundant structures, accounting for >90% coverage, are represented for both PD-1 and PD-1 NF proteins. Glycan composition is indicated with the convention "HNFS", which represents the number of N-acetyl-hexosamine (H), hexose (N), sialic acid (S) and fucose (F). The most abundant glycan species contained in the two proteins have comparable compositions that differ only for one fucose unit. Related glycan species are connected by lines.

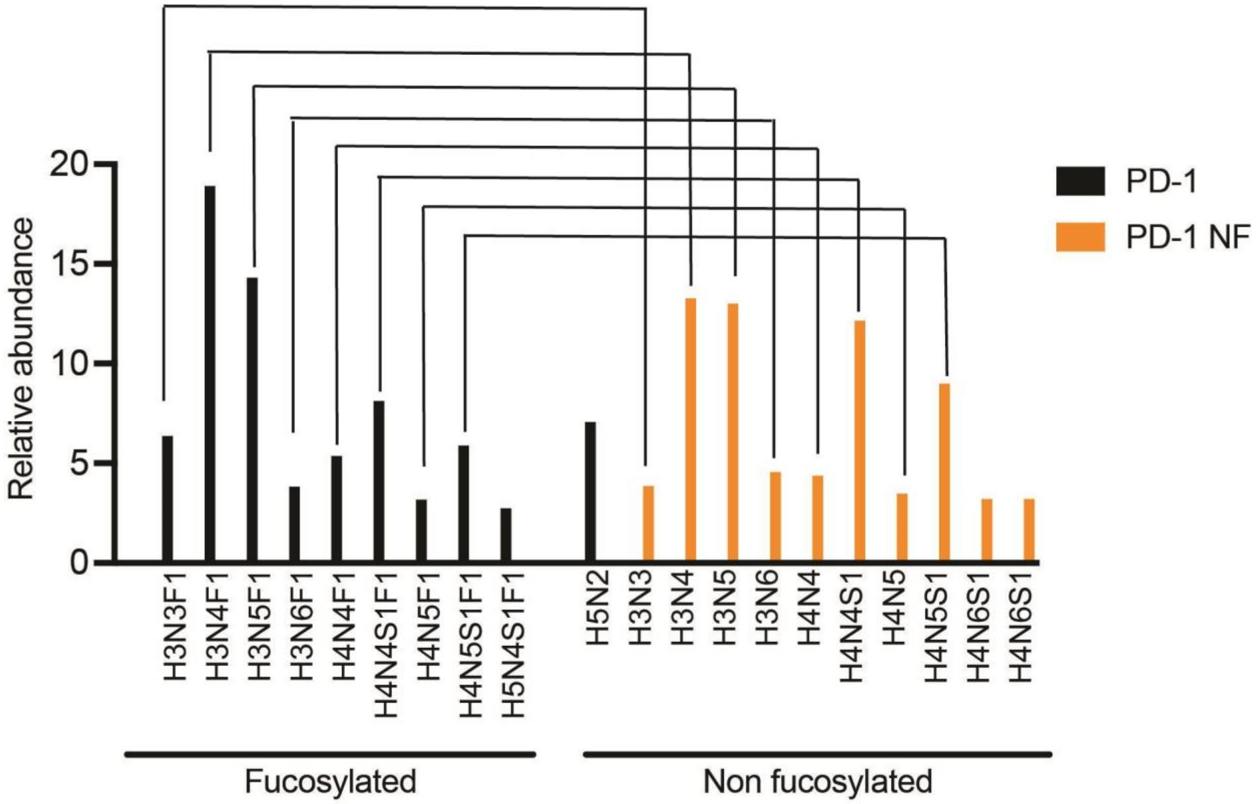

**Supplementary Figure 5. Representative sensogram for determination of the binding kinetics of anti-PD-1 antibodies.**

A3 to G3 are samples containing PD-1 at different concentrations (from high to low); H3 is buffer.

The sequential steps separated by a dotted line represent:

1. Baseline in assay buffer
2. Baseline in PD-1 analyte to detect potential nonspecific interactions
3. Baseline in assay buffer
4. Antibody capture
5. Baseline in assay buffer
6. Association with PD-1 protein in different concentrations (A-G)
7. Dissociation in assay buffer

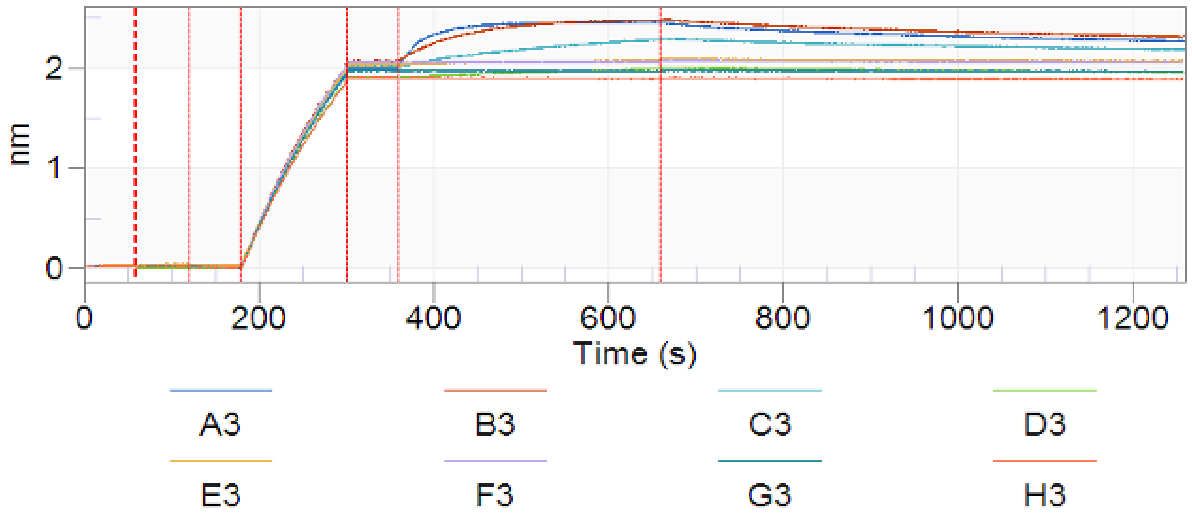

**Supplementary Figure 6. Core fucose on the PD-1 N-glycans does not modulate binding to PD-L1.** Flow cytometry analysis of CHO-K1 cells or CHO-K1 *Fut8* KO cells transfected with full length PD-1 and stained with PD-L1-Fc protein. The graphs indicate average values of n=2.

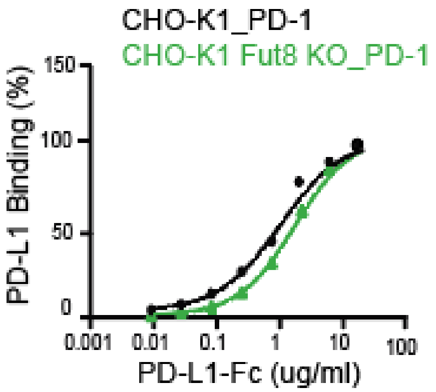

**Supplementary Figure 7. Fucose inhibitors efficiently reduce cell surface fucosylation and do not impact cell viability. A.** Flow cytometry analysis of Jurkat cells treated with 2F-Peracetyl-Fucose in concentrations ranging from 0 uM to 600uM. Fucose content was determined by staining with AAL or LCA lectin. **B.** Cells treated with 0 uM, 50 uM, 100 uM, 300 uM, 600 uM of 2F-Peracetyl-Fucose (from left to right) were stained with Zombie red viability dye to determine the percentage of the viable cells.

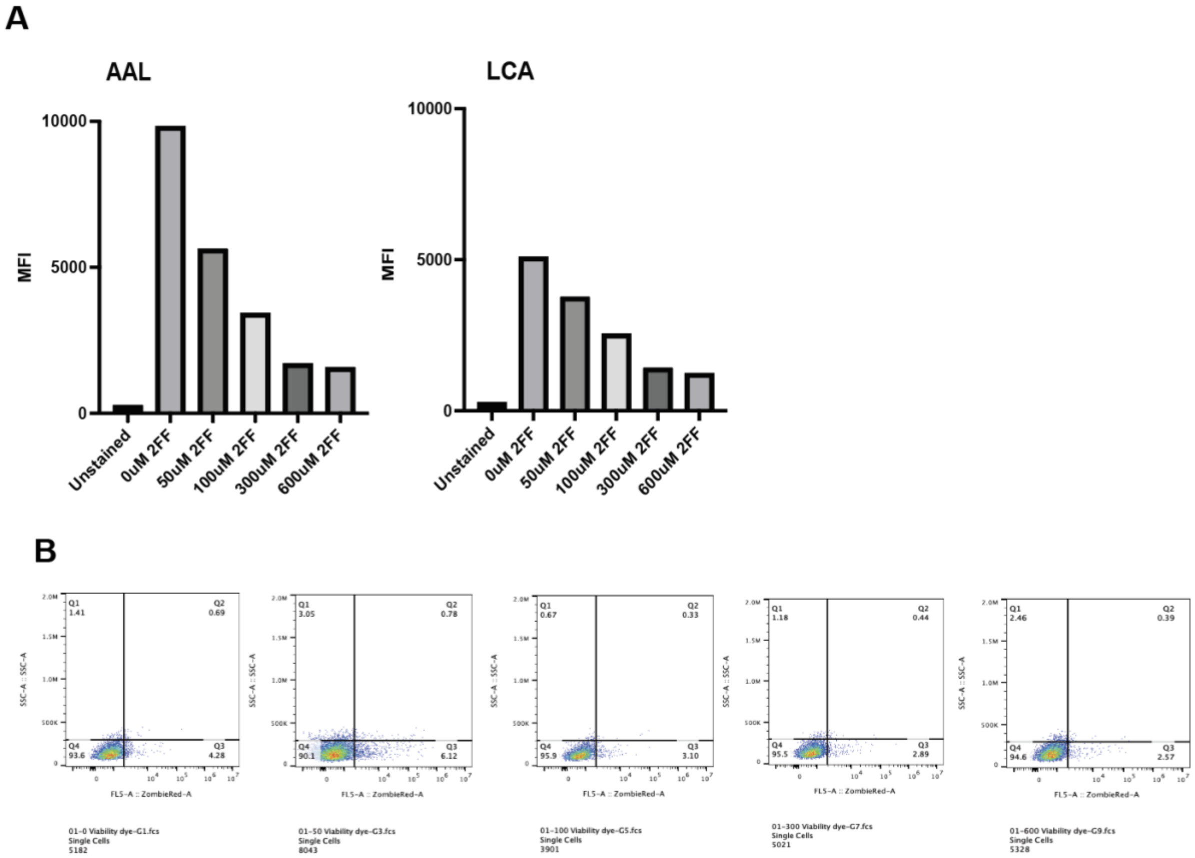

**Supplementary Figure 8. Quantification of distinct PD-1 variants by different plate-based assays.** Combination of antibodies for coating and detection allows the analysis of total or non-fucosylated PD-1. Human PD-1 Quantikine ELISA preferentially detects PD-1 with core fucose at N58. Human PD-1 DuoSet ELISA uses polyclonal antibodies and detects PD-1 independently from glycosylation content.

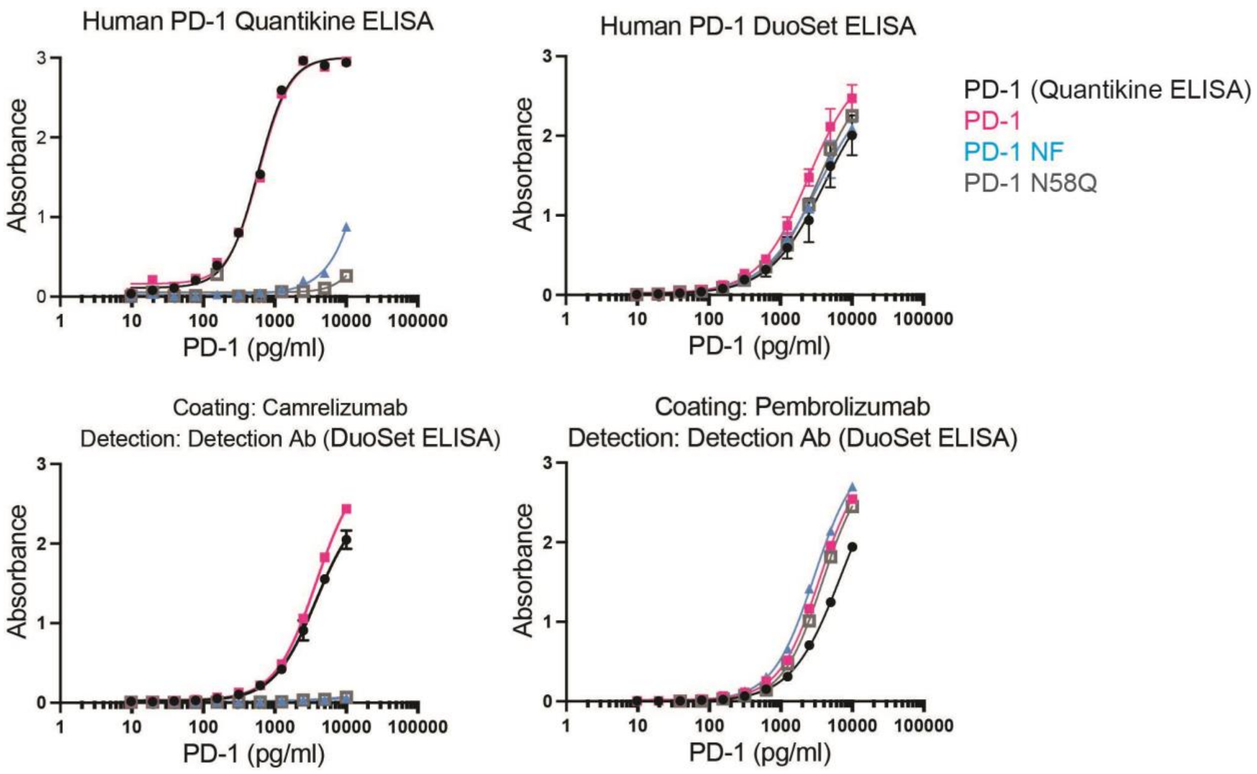
